# Large single-locus effects for maturation timing are mediated via body condition in Atlantic salmon

**DOI:** 10.1101/780437

**Authors:** Paul V Debes, Nikolai Piavchenko, Annukka Ruokolainen, Outi Ovaskainen, Jacqueline E Moustakas-Verho, Noora Parre, Tutku Aykanat, Jaakko Erkinaro, Craig R Primmer

## Abstract

Maturation timing is a pivotal life-history trait balancing probabilities between mortality and reproduction. Environmental vs. genetic contributions to traits associated with maturation initiation, such as growth and body condition, remain uncertain because of difficulties in determining causality. In Atlantic salmon, maturation timing associates with a large-effect locus around *vgll3*, but how this locus affects maturation remains unknown. We combined controlled breeding with common-garden experimentation at two temperatures and show that *vgll3* effects on maturation of males express primarily via body condition, which we demonstrate in the males’ non-maturing female relatives, thus avoiding reverse causality. Between homozygous *vgll3* genotypes, maturation probability differed several folds and female condition differed 2% in both temperature environments. *Vgll3* effects explained 25 and 16% of maturation probability heritability and 15 and 6% of female condition heritability, in the warm and cold environment, respectively. Non-significant *vgll3* effects on female length were antagonistic to those on condition but of equal proportional size. When controlling for *vgll3* effects, genetic correlations changed antagonistically between both maturation and condition vs. growth, suggesting *vgll3* as a resource-allocation locus. The results support large *vgll3* maturation effects being mediated via environmentally stable body condition effects, enabling rapid co-evolution between the life-history traits.

## Introduction

Sexual maturation is a central life-history trait that contributes to maximizing individual survival and reproductive success and, thereby, per *capita* population growth rate (Stearns 1992; Roff 2002; Wells et al. 2017). Maturation initiation is assumed to be genetically and environmentally controlled via factors including growth or size and body condition (Stearns 1992; Roff 2002; Taranger et al. 2010; Wells et al. 2017; Andersson et al. 2018), which may signal current energy status (Danielsen et al. 2013; Dupont et al. 2014). However, disentangling cause and effect is a major challenge in studies on maturation and associated traits (Kause et al. 2003; Cousminer et al. 2013; Bell et al. 2018). For example, as opposed to growth controlling maturation, ongoing maturation can temporally boost growth, such as the human puberty growth spurt. Likewise, the maturation process can reduce somatic growth by competing with resources and lower condition by depleting reserves, or affect both growth and condition via appetite (Stearns and Koella 1986; Taranger et al. 2010; Andersson et al. 2018). Related to this problem, fundamental evolutionary knowledge on presence and relative importance of genetic vs. environmental contributions to maturation timing and their link to maturation component traits, such as growth or condition, remains limited (Gjedrem and Baranski 2005; Law 2007; Dunlop et al. 2009; Enberg et al. 2012).

Contrary to the assumption that life-history traits are comprised of several underlying component traits with polygenic architecture (Lande 1982; Roff 2002), each coded by many loci with small effect on phenotypic variation (Lynch and Walsh 1998; Hill 2010), maturation timing in Atlantic salmon (*Salmo salar*) associates with a locus explaining a large proportion of phenotypic variation (33-39%; Ayllon et al. 2015; Barson et al. 2015). This finding bears implications on evolutionary predictions because the expected selection response differs depending on whether a trait is governed by only small- or also large-effect loci (Roff 2002; Kuparinen and Hutchings 2017). For example, initial allele frequencies for the latter, but not the former, may govern evolutionary responses (Kardos and Luikart 2019). The large-effect locus also offers outstanding opportunities for understanding the role of age-specific body size and condition and for quantifying the relative importance of genetic vs. environmental contributions of these traits to maturation timing. The large-effect locus positions close to a transcriptional co-factor gene (*vgll3*), hypothesized as a strong candidate gene (Ayllon et al. 2015; Barson et al. 2015). Known functions of Vgll3, such as inhibition of adipocyte differentiation in cell lines in favor of somatic growth processes and its negative transcriptional correlation with body mass and fat content in mice (Halperin et al. 2013), suggest a mechanistic link of *vgll3* with maturation via control of resource allocation between energy reserves and somatic growth. Genetic markers in the genome region around *vgll3* have also been associated with salmon length (Barson et al. 2015), and human maturation and growth (Cousminer et al. 2013), body condition (Nakayama et al. 2017), or condition change during maturation (Tu et al. 2015), fostering the idea that the link between this locus and maturation, somatic growth, and condition underlies a common functional phenotype. However, a comprehensive joint assessment of independent *vgll3* effects on maturation, growth, and condition, which also requires that the latter two traits are estimated free of maturation-induced changes, is still missing.

In Atlantic salmon, sexual maturation studies have a long history (Meerburg 1986; Marschall et al. 1998; Andersson et al. 2018) and this species offers features allowing for a joint assessment of maturation, growth, and condition, thus enabling independent assessment of their links with *vgll3*. Specifically, readily available pedigreed hatchery populations allow for planned breeding of highly fecund individuals. Many offspring with tightly connected pedigrees combined with common garden experimentation followed by quantitative genetic analyses enable i) dissection of genetic from environmental effects, ii) estimation of genetic correlations between environments or iii) estimation of genetic, environmental, and phenotypic correlations between traits. Perhaps the biggest advantage, however, is the observation that Atlantic salmon males, but usually not females, can mature during their first year (Marschall et al. 1998). This provides opportunities to estimate environmental and genetic contributions, including those of *vgll3*, to maturation in males and to growth and condition in non-maturing females. By genetically relating male maturation to maturation-component traits of their female relatives, it is possible to infer the presence and relative importance of genetic vs. environmental components of maturation timing and its maturation-unbiased component traits.

Here, we implemented a quantitative genetic breeding design for 42 pedigreed Atlantic salmon parents with known homozygous *vgll3* genotypes and studied >5,000 male and female offspring from 41 families with all *vgll3* genotypes (**figure 1A, B**). The study of these families in a common-garden environment enabled general estimates of genetic vs. environmental contributions for maturation timing of first-year Atlantic salmon and its component traits in conjunction with an assessment of specific *vgll3* effects on maturation, growth, and condition. Further, we evaluated whether a known association of the *vgll3* locus with maturation (co)expresses via growth, condition, or both. We did so by testing whether male maturation probability correlates genetically with maturation-unbiased female somatic growth or body condition (or with both) and by quantifying the *vgll3* contribution to the genetic between trait correlation between sexes. By replicating all families in two environments with a life-time 2 °C water-temperature difference across a seasonal temperature curve (a likely global-warming scenario (IPCC 2014); **figure 1C, D**), we were also able to assess environmental effects and their influences of wide relevance to our estimates. We show that large *vgll3* effects on maturation timing are mediated via large *vgll3* effects on body condition and that allele specific phenotypic effects are environmentally stable, although the relative contributions to the phenotypic variance differ with water temperature. Combined, our results also suggest *vgll3* as a candidate locus for controlling the resource allocation trade-off between energy reserves and somatic growth.

**Figure 1:**
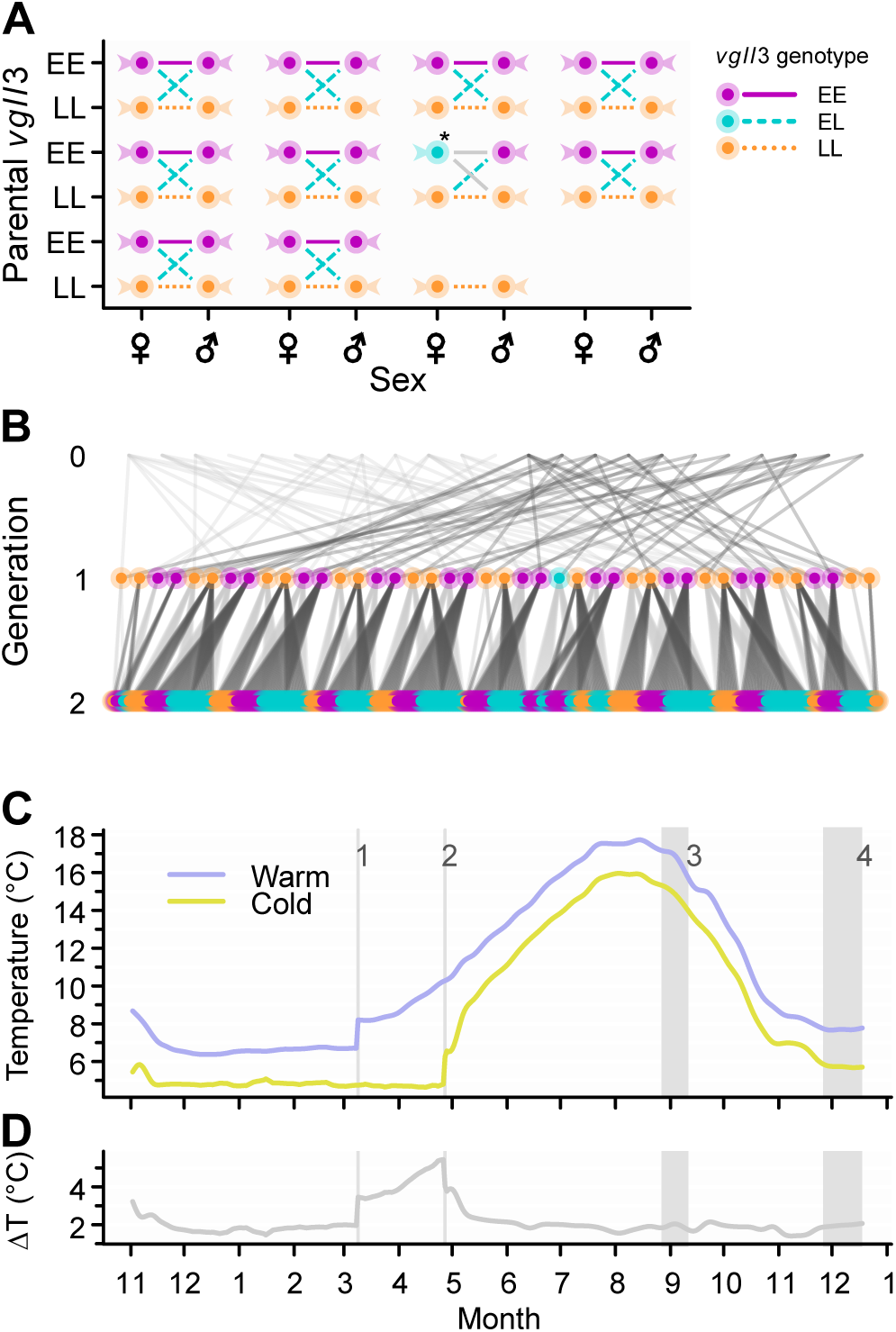
Breeding design for *vgll3*, pedigree, and water temperatures during the experiment. (**A**) Realized breeding design with anticipated 2×2 factorial for 24 *vgll3* homozygous dams by 24 *vgll3* homozygous sires that was successful for nine factorials. One factorial curtailed because of low egg survival, whereas one dam (marked by asterisk) was determined to be *vgll3* heterozygous upon confirmatory genotyping after spawning. (**B**) Pedigree depicting relationships among studied salmon (generation 2; N = 5,145; only every 10^th^ individual plotted to facilitate viewing) via parents (generation 1; N = 42) up to their grandparents (generation 0; N = 23). Upward links to dams and sires in the pedigree are colored in light and dark gray, respectively. *Vgll3* genotypes for created families (lines, **A**) and breeders (fish shapes, **A**), or in generations 1 and 2 (circles, **B**), are depicted corresponding to the legend next to **A**. (**C**) Average temperature across eight tanks in the warm or cold temperature environments. Experimental events are indicated by light gray vertical lines with 1: transfer from egg incubators (N = 2) to tanks (N = 8) and first feeding in the warm environment; 2: transfer from egg incubators (N = 2) to tanks (N = 8) and first feeding in the cold environment; 3: phenotyping of length and condition as used in analyses described in the manuscript; 4: maturation status and sex confirmation assessments. (**D**) Average temperature differences between the warm and cold temperature environments resulting from **C**.

## Methods

### Fish population, breeding and experimental design, data collection

The experimental cohort was parented by pedigreed hatchery fish maintained by the Natural Resources Institute Finland (Laukaa, Finland). The hatchery-stock ancestors originated from the River Neva, Russia, which drains into the Baltic Sea. In November 2017, we crossed 48 parents with known *vgll3* genotype as 12 2×2 factorials of unrelated *vgll3* homozygous individuals; each factorial yielded four reciprocal *vgll3* offspring genotypes (EE, EL, LE, LL; details on realized design and resulting pedigree in **figure 1A, B**). We reared the experimental cohort in two recirculation systems controlled for water temperature, oxygen, dissolved nitrate components, and natural light cycle, which affect growth or sexual maturation timing (Taranger et al. 2010; Andersson et al. 2018). We split each family by randomizing ∼400 eggs across four egg-incubator replicates in two water temperatures (kept in darkness). At first feeding, we randomized an equal number of individuals from each incubator into eight similar tank replicates for each temperature (due to differential egg mortality: mean = 5, range: 1-7, total per family across 16 tanks: mean = 155, range: 21-234). Due to mortality, removal of sick individuals, unknown genotype or sex, culling of random individuals to reduce densities, and removal of individuals with mismatch between genotypic and phenotypic sex (see below), an average of 125 individuals (range: 11-199) per family existed for analysis. Water temperatures in egg incubators and tanks followed a seasonal cycle with a 2 °C difference, referred to as “warm” or “cold” (warm, range: 6.3-17.7 °C; cold, range: 4.1-16.0 °C; **figure 1C, D**). Fish were fed *ad libitum* using a commercial salmon diet starting 2018-03-09 (warm) or 2018-04-27 (cold) (**figure 1C**). Once fish size allowed passive integrated transponder tagging (warm: August 2019; cold: September 2019) to enable re-identification, we anesthetized (using methanesulfonate), fin clipped, and tagged individuals; fin clips allowed for genotyping individuals to assign family, determine molecular sex, and confirm *vgll3* genotype. We measured fork length (± 1 mm) and weighed wet mass (± 0.01 g) in September (the earliest time point when all fish were individually tagged), and in December (at final spawning time) when we also determined maturity status by checking for extruding milt while gently pressing the abdomen (**figure 1C**). During December sampling, we culled and dissected 84% of all fish (N = 4,318; others were kept for later studies). Genotypic and phenotypic sex matched in 99.9% of the culled fish (five culled fish, i.e., 0.1%, diverged between genotypic and phenotypic sex and were removed from analysis), and we used phenotypic sex in 109 fish with unknown genotypic sex. We confirmed maturity status in 100% of the culled fish. Once individual identification was possible, we applied a feed-restriction treatment (either *ad libitum* feeding for seven days per week or *ad libitum* feeding for two days per week with no feeding for two or three days between feedings) that was crossed with the temperature treatment for a five-week period (September - October 2018), but we did not detect any feed-restriction effect on maturation (**figure A1A**, appendix). Because length and condition data analyzed originated from before the feed restriction, we omitted the feed restriction term from further analyses. Animal experimentation was conducted according to European Union Directive 2010/63/EU and under license ESAVI-2778-2018 by the Animal Experiment Board in Finland (ELLA).

Genotypes and genotypic sex of both parents and the experimental cohort were determined using a multiplex-PCR for 177 single nucleotide polymorphisms (SNPs) of a previously described SNP panel (Aykanat et al. 2016), followed by Ion Torrent (984 broodstock individuals from which we drew parental individuals) or Illumina platform (MiSeq or Next-Seq) sequencing (parental individuals, experimental offspring cohort). Using a subset of unlinked, polymorphic SNPs, we reconstructed the experimental cohort’s grandparents, i.e., parents of the used parents (with 131 SNPs), under maximum likelihood (Jones and Wang 2010) (“Settings for the parental reconstruction” in the appendix, available online). and we assigned the experimental individuals to their 48 parents (with 141 SNPs) with a likelihood approach (Anderson 2010). Merged information about reconstructed grandparents and assigned parents yielded a three-generation pedigree (**figure 1B**) on which we based the relationship matrix utilized in animal-model analyses (Henderson 1973).

### Data analysis

We fitted a series of uni- and multivariate general and generalized linear animal models to data for maturation binaries recorded in December and for length and condition recorded in September. We defined condition as deviation from the (temperature-specific) slope of logarithmic mass on logarithmic length - a correlate of first-year salmon lipid content (Sutton et al. 2000). Length and condition data were first log-transformed, then mean-centered and variance scaled (across temperatures) to allow for estimating biologically meaningful proportional and phenotypic variance-standardized effects.

To determine the multivariate model structure and allow for comparisons between uni- or bivariate with multivariate model estimates (**figure A2**, appendix), we first fit univariate models for the binary response of male maturation and bivariate models for the continuous responses of sex-specific length or condition. We estimated models effects for the binary maturation response (including the multivariate model) using Bayesian Markov Chain Monte Carlo simulations, which is recommended for binary animal models (de Villemereuil et al. 2013), with the R-package MCMCglmm v. 2.28 (Hadfield 2010) in R v. 3.5.3. For the continuous responses, we initially fitted bivariate models under REML, which is recommended for continuous animal models (de Villemereuil et al. 2013), using ASReml-R v. 3 (Butler et al. 2009) in R v. 3.0.2. For Bayesian models, we removed mean and variance terms when all estimates included zero in their 95% credible intervals (see below; **figures A1, A3**, appendix). For REML models, we removed fixed terms with *P* ≥ 0.05 for Wald’s *F*-test with denominator degrees corrected for small samples (Kenward and Roger 1997) and removed random terms with *P* ≥ 0.1 for likelihood ratio tests if parameters were restricted to be ≥ 0 and with *P* < 0.05 otherwise (Stram and Lee 1994) (see below; **table A1**, appendix). We then fitted the multivariate model corresponding to the chosen univariate (maturation) or bivariate models (sex-specific length or condition), and by adding the required between-trait covariances for the additive genetic effects (i.e., 2,599 [males] or 5,209 [both sexes] relationship-matrix-predicted breeding values) (Henderson 1973), common environmental effects (i.e., 16 tanks), maternal effects (i.e., 21 dams), and individual environmental effects including measurement error and non-additive genetic effects (i.e., 2534 [males] or 5145 [both sexes] residuals). We fitted the models with probit-link function and residual variance fixed to one for maturation, corresponding to genetic threshold models (Hadfield and O’Hara 2015), and with identity link function for length and condition.

We did not detect maternal effects on any trait (**figure A1B, table A1**, appendix; we removed dam effects as a consequence) and no common environmental effects on maturation (**figure A1B**, appendix; we still kept the experimental tank effects), but on length and condition, which contributed up to 6 and 15% to the total phenotypic variance, respectively (**figure A2**, appendix). We started model selection with mean-effect interactions and removed non-significant effects (**figures 2; A1A, A3**, appendix). Final models followed the general equation (per trait), with colon indicating term interaction and variance terms in italic: y ∼ Intercept + Temp + Vgll3 + *Temp:Tank* + *Temp:Animal* + *Temp:Residual*. The Vgll3 model term refers to either three genotypes (reciprocal heterozygote differences were absent; posterior EL-LE contrast, 95% CI: MAT_EL_-MAT_LE_ = 0.00, −0.47-0.49), or to the additive allelic effect (we fitted both; see below); the Temp model term refers to a factor for warm or cold environments. Covariance matrices across temperatures for initially fit univariate models were diagonal for tank and residual effects and unstructured for dam and animal effects. The latter allows estimating dam- and additive genotype-by-environment effects (Falconer 1952). For the multivariate models (five response traits: sex-specific length, sex-specific condition, male maturation) we expanded the covariance matrices to temperature-specific block-diagonals for *Trait:Temperature:Tank* (two 5×5 blocks), a full *Trait:Temperature:Animal* genetic covariance matrix (10×10), and temperature- and sex-specific block-diagonals for *Trait:Temperature:Residual* (two 2×2 female and two 3×3 male blocks; **files A1-3**, electronic enhancements). We used recommended (co)variance priors following a 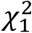 distribution in Bayesian univariate threshold models (de Villemereuil et al. 2013) or priors that resulted in flat priors for heritabilities and correlations in multivariate models. We based estimates on 5,000 or 1,000 retained iterations after 50,000 burnin iterations and sampling every 1,250 and 2,500 or 3,000 iterations (uni- and multivariate models, respectively). We invoke latent variable truncation between −7 and 7 to avoid numerical errors. We confirmed model convergence by trace plot inspection and ensuring lag-two autocorrelation < 0.1 for all estimates.

**Figure 2:**
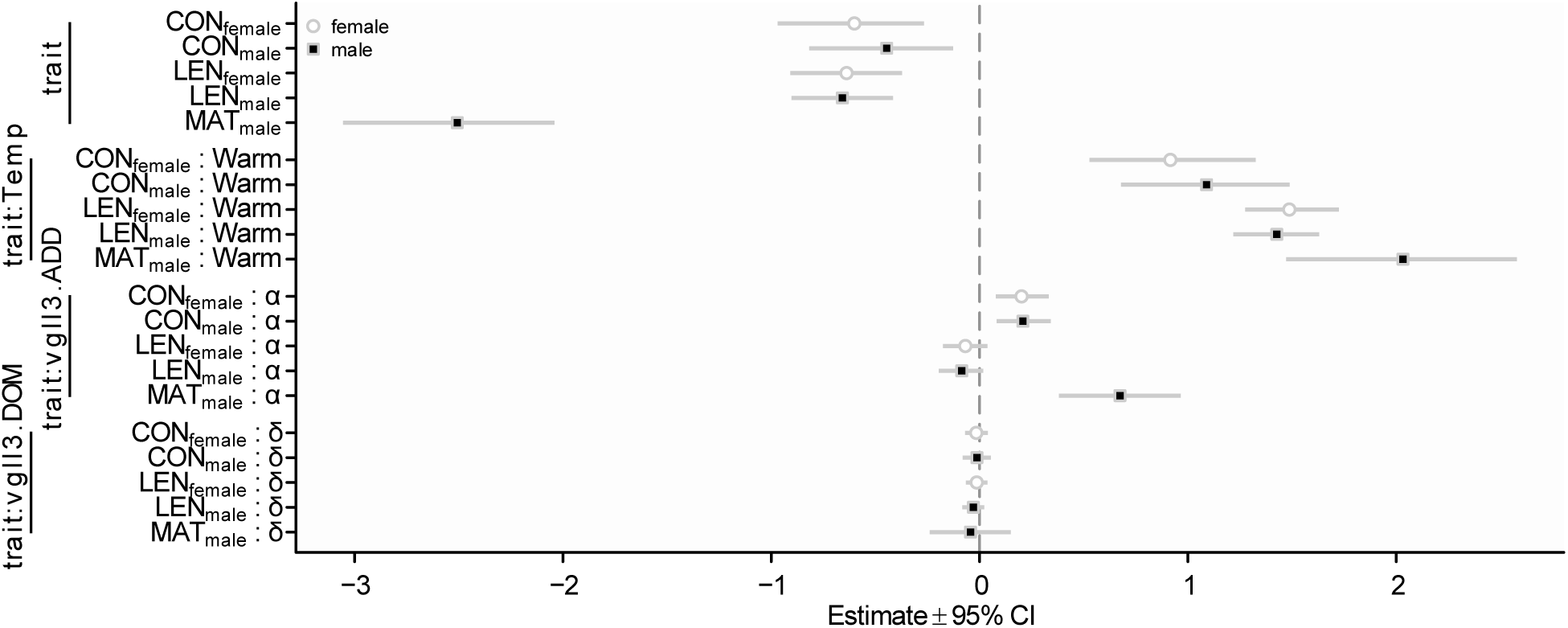
*Vgll3* dominant effects are absent. Mean and contrast effect estimates (grouped by term, with term level indicated) with 95% credible intervals for the multivariate animal model on traits of male maturation (MAT_male_; probit scale), male length (LEN_male_; standardized log of cm), female length (LEN_female_; standardized log of cm), male condition (CON_male_; standardized log of g, adjusted to common size), and female condition (CON_female_; standardized log of g, adjusted to common size). Traits were modelled as a linear function of temperature effects (Temp; levels: Cold, Warm), additive *vgll3* effects (vgll3.ADD), and dominant *vgll3* effects (vgll3.DOM). Dominant *vgll3* effects for all traits were removed because their 95% credible intervals included zero. All effects estimates originate from a common multivariate generalized animal model (N_families_ = 41, N_male_ = 2,534, N_female_ = 2,611).

We tested and estimated additive and dominant *vgll3* effects by replacing the factorial Vgll3 term by appropriate covariates (α: −1, 0, 1; δ = 0, 1, 0). To obtain observed-scale maturation probability parameter estimates, we integrated over marginal model predictions and used methodology implemented in the R-package QGglmm v. 0.7.2 (de Villemereuil et al. 2016) in R v. 3.5.3, applied to each retained MCMC iteration to estimate credible intervals. We calculated heritability as the ratio of additive genetic over total phenotypic variance, and estimated genetic, environmental, and phenotypic correlations following Searle (1961). All necessary data and an R-script mirroring the final uni- and multivariate modeling have been deposited in the Dryad Digital Repository (during review: https://datadryad.org/stash/share/FLBZRu6qosNz909duxMd6Tw-LeGo16cPHlfprRaqKb0; after acceptance: https://doi.org/10.5061/dryad.jh9w0vt6k).

## Results and Discussion

We used a multivariate generalized animal model to test for associations between *vgll3* and each of the investigated traits of male maturation probability (maturation, MAT), sex-specific somatic growth (length, LEN), and sex-specific body condition (condition, CON). To avoid reverse causality (i.e., inferring condition- or length-mediated *vgll3* effects on maturation that are truly maturation-mediated *vgll3* effects on somatic growth or condition) we focused on *vgll3* effects on maturation in males and on *vgll3* effects on length and condition in females (of which none matured). To allow for wider interpretation and relevance of our results, we report maturation results, consistent with a genetic threshold model, on the probit or observational probability scale, which are relevant for breeding-value based or phenotypic selection, respectively (de Villemereuil et al. 2016). Likewise, we report results for length and condition on the proportional scale, which is relevant to growth processes at the individual level, or on the standardized effects size scale (phenotypic standard deviation across environments, psd), which is relevant among individuals at the population level.

### *Vgll3* dominance effects are absent

We first formally tested jointly for additive (α) and dominant (δ) effects for all traits, because sex-specific dominance (i.e., *vgll3*-by-sex epistasis) has been reported for sea age at maturity in two of three previous studies (Barson et al. 2015; Czorlich et al. 2018; but see Ayllon et al. 2019). When fully accounting for experimental randomization and relatedness, we did not detect dominance for any trait (**figure 2**). This result suggests that previous inferences about *vgll3*E* dominance on maturation age in males from wild populations (Barson et al. 2015; Czorlich et al. 2018) cannot be generalized to all situations. However, dominance absence verifies a requirement for meaningfully associating *vgll3* effects between male and female traits that would not be fulfilled under a sex-by-locus interaction, such as sex-specific dominance, although we cannot exclude the presence of non-additive allelic effects on prospective female maturation, which, however, does not affect inferences for any trait in the present study.

### *Vgll3* associates with maturation timing and body condition across environments

After removing dominance effects, we detected *vgll3* effects on maturation probability and, in both sexes, on condition, but not on length (i.e., the 95% credible intervals included zero), in both temperature environments; the *vgll3*\**E* allele associated with both higher maturation probability and higher condition (**figure 3**). These results provide experimental confirmation that the allele associated with earlier maturation following marine migration (Ayllon et al. 2015; Barson et al. 2015; Ayllon et al. 2019) has consistent effects in males maturing in freshwater, and that the same allele also associates with a higher condition. All focal trait means were much higher in the warm than in the cold environment (**figure 3;** posterior temperature contrasts, 95% credible interval [95% CI]: probit scale, MAT_Warm_ - MAT_Cold_ = 2.0, 1.5-2.6; psd scale, female LEN_Warm_ - LEN_Cold_ = 1.5, 1.3-1.7; female CON_Warm_ - CON_Cold_ = 0.92, 0.57-1.29). Despite these strong environmental effects on trait means (discussed below), *vgll3* effect estimates did not differ between temperature environments for any trait (**figure 3; figure A3**, appendix). For male maturation probability, we estimated a positive additive *vgll3*\*E allele effect on the probit scale that translated to considerable maturation probability differences among *vgll3* genotypes (**figure 3A, B)**. For female length, we estimated a non-significant and negative additive *vgll3*\*E allele effect on the psd scale of α = −0.06 (95% CI = −0.16-0.05). This effect translates to a non-significant 1% lower body length per *vgll3*\*E allele (**figure 3C, D**). For female condition, we estimated an additive *vgll3*\*E allele effect on the psd scale of α = 0.20 (95% CI = 0.09-0.33). This among-individual additive *vgll3* effect on condition is much larger than that estimated for length. However, it translates to the same proportional effect as estimated for length, but in the opposite direction. Specifically, a non-significant 1% body length decrease per *vgll3*\*E allele is accompanied by a significant 1% body mass increase (**figure 3E, F**). Although the *vgll3* effect on length is non-significant and of little, if any, importance at the population level (but may still represent a within-individual trade off, discussed below), the same effect size of 1% body mass (condition) increase per *vgll3*\*E allele is very likely relevant for maturation. Specifically, male salmon gonad development is dependent on body reserves, such as fat (Taranger et al. 2010; Andersson et al. 2018), and gonads constitute 5-12% of body mass at spawning for the investigated freshwater life stage (Rowe et al. 1991; Fleming 1996; Trombley et al. 2014). Thus, ignoring unknown reserve-to-gonad mass conversion factors, condition variation explained by *vgll3* genotype may finance up to 40% of the required extra mass (in *vgll3* EE vs. LL; assuming minimum gonad proportion; 2% / 5% = 0.4;). These results provide the first indication that the large effect of *vgll3* on maturation is, at least partly, mediated by a large effect of *vgll3* on condition. We strengthen this further by providing additional evidence below where we quantify *vgll3* effects on trait-specific heritabilities and between-trait genetic correlations.

**Figure 3:**
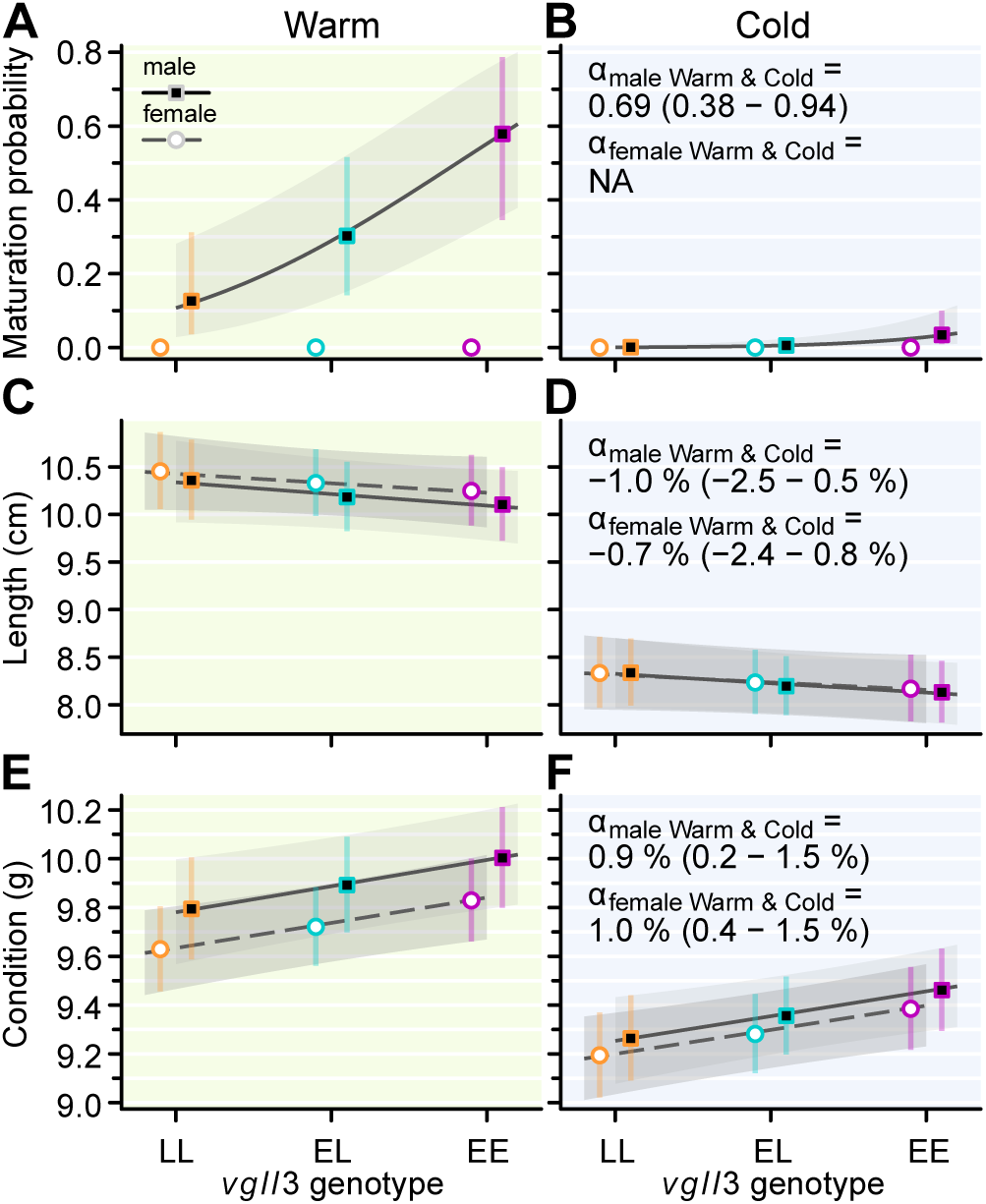
*Vgll3* associates with maturation and body condition across environments. (**A-F**) Back-transformed mean and additive effect (α) estimates with 95% credible interval per *vgll3* genotype for male maturation probability (**A, B**; α on probit scale; no female maturation occurred), sex-specific body length (**C, D**) sex-specific body condition (**E, F**; standardized mass for a geometric mean-sized fish of 9.1 cm). Mean effect estimates are valid for either a warm (**A, C, E**) or a 2 °C colder (**B, D, F**) environment. All *vgll3* genotype mean or additive effects estimates originate from common multivariate generalized animal models (one for mean effects, one for additive effects; N_families_ = 41, N_male_ = 2,534, N_female_ = 2,611). Additive *vgll3* effect estimates with 95% credible intervals (**B, D, F**) are valid for both environments (see also **figure A3**, appendix).

### *Vgll3* explains heritability for maturation timing and body condition, and the genetic between-trait correlation between sexes

The same multivariate models also enabled us to estimate environment- and trait-specific heritabilities (h^2^) and trait-specific genetic correlations (R_G_) between temperature environments (both contribute to genotype-by-environment interactions; Sae-Lim et al. 2016) and also to estimate how much *vgll3* contributes to either. The heritability of a trait quantifies the proportion of phenotypic variance explained by the sum of all underlying additive genetic effects and predicts the response to phenotypic selection. The genetic between-environment correlation quantifies the genotype re-ranking between environments and predicts, in conjunction with heritability estimates, the response to selection across environments (Falconer 1952). Fitting each model with and without specified additive *vgll3* effects, enabled us to quantify the relative genetic and phenotypic contributions of additive genetic *vgll3* effects, and thus the trait-specific evolutionary importance of *vgll3* within and across environments. It should be noted that significance of *vgll3* effects is tested via the above-reported phenotypic effects, not via heritability differences when *vgll3* effects contribute or not.

Remarkably, male maturation heritability (h^2^MAT) on both the probit and the observed scale was 1.4 times larger in the warm environment than in the 2 °C colder environment (**figure 4A, B; file A4**, electronic enhancement; probit scale, posterior temperature contrast, 95% CI; h^2^MAT_Warm_ - h^2^MAT_Cold_ = 0.32, 0.21-0.41; observed scale, posterior mean, 95% CI; h^2^MAT_Warm_ = 0.41, 0.32-0.50; h^2^MAT_Cold_ = 0.10, 0.04-0.16; posterior temperature contrast, 95% CI; h^2^MAT_Warm_ - h^2^MAT_Cold_ = 0.28, 0.17-0.38;). In addition to heritability differences, the between-temperature genetic correlation estimate for maturation (R_G_MAT_Warm_._Cold_) was < 0.8 (posterior mean, 95% CI; R_G_MAT_Warm_._Cold_ = 0.70, 0.45-0.93), which is somewhat lower than most previous genotype-by-temperature environment estimates for maturation of related species (Sae-Lim et al. 2016). Importantly, R_G_ < 0.8 indicates breeding value re-ranking between environments with a mere 2 °C temperature difference and thus is of relevance to natural and aquaculture settings (Robertson 1959; but see Sae-Lim et al. 2016). This genetic correlation estimate is of particular relevance to predicting selection response across environments in combination with the pronounced environmental differences in maturation heritability (Falconer 1952), and the values reported here readily apply to a 2 °C global-warming scenario (IPCC 2014). For maturation, additive *vgll3* effects on the probit scale explained 24.8 and 16.1% of heritability in the warm and cold environments, which explains (by multiplication with *vgll3*-unadjusted heritability) 16.7 and 7.5% of phenotypic variance, respectively. On the observed-scale, *vgll3* effects explained 32.9 and 28.1% of heritability and 13.6 and 2.7% of phenotypic variance, respectively. Additive *vgll3* effects also considerably reduced the between-environment genetic correlation for maturation (**figure 4C; file A4**, electronic enhancement), which supports an interpretation that the abovementioned result for non-significant temperature-by-*vgll3* interaction effects imply environmentally stable *vgll3* effects relative to other, less environmentally stable additive genetic effects on maturation.

**Figure 4:**
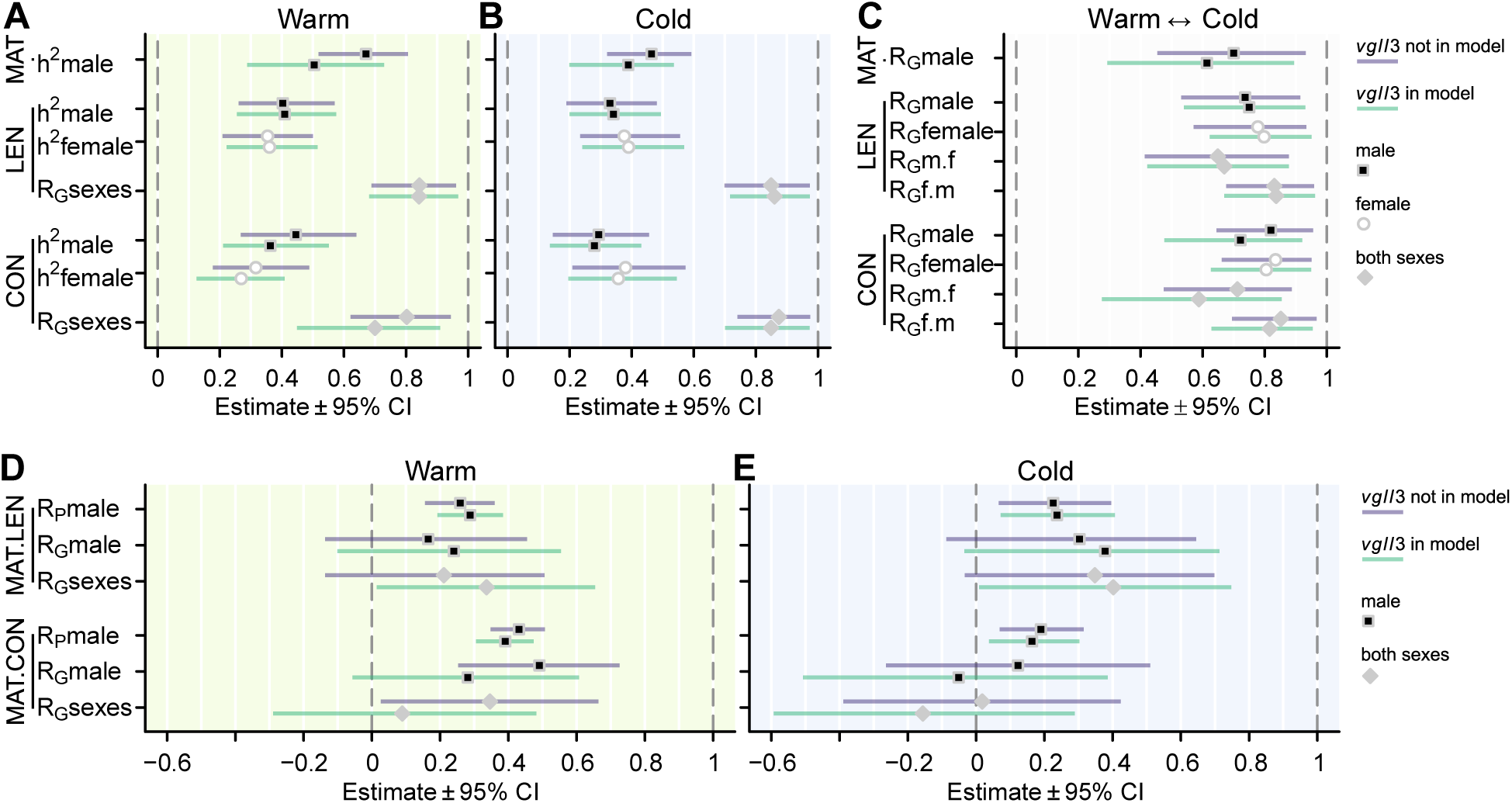
*Vgll3* explains heritabilities and genetic between-trait correlations. (**A, B**) Environment- and sex-specific heritability (h^2^) and between-sex genetic correlation (R_G_sexes) estimates with 95% credible intervals for male maturation (MAT), sex-specific body length (LEN), and sex-specific body condition (CON). Estimates are valid for either a warm (**A**) or 2 °C colder (**B**) environment. (**C**) Between-environment genetic correlation (R_G_) estimates with 95% credible interval within sex (R_G_male, R_G_female) and reciprocally between sexes (R_G_m.f, R_G_f.m) for male maturation (MAT), sex-specific length (LEN), and for sex-specific condition (CON). Estimates refer to the correlation between a warm and a 2 °C colder environment. (**D, E**) Between-trait phenotypic correlation (R_P_) estimates with 95% credible intervals between male maturation and male length (MAT.LEN) or male maturation and male condition (MAT.CON) and between-trait genetic correlation (R_G_) estimates with 95% credible intervals between male maturation and sex-specific length (MAT.LEN) or male maturation and sex-specific condition (MAT.CON). Estimates are valid for either a warm (**D**) or 2 °C colder (**E**) environment. All estimates were obtained by two similar multivariate models (N_families_ = 41, N_male_ = 2,534, N_female_ = 2,611), differing in whether additive *vgll3* additive effects were included (the same model underlying estimates in **figure 3**) or excluded as indicated in the legends.

Female length heritability was, in contrast to that for maturation probability, similar between environments (posterior mean, 95% CI; h^2^LEN_Warm_ - h^2^LEN_Cold_ = −0.03, −0.19-0.13) and the between-environment genetic correlation of R_G_LEN_Warm_._Cold_ = 0.78 (0.57-0.93) indicated less genotype re-ranking between environments than that for male maturation (**figure 4A-C**). This between-environment genetic correlation contrasts with much lower estimates in other fish species (Sae-Lim et al. 2016). Accounting for *vgll3* effects, unexpectedly, increased female length heritability, as opposed to lowering it as observed for maturation heritability, though by only 1.5 and 3.5% in the warm and cold, respectively (**figure 4A, B)**. When we investigated underlying variance components between models for length, we found that including *vgll3* effects re-allocated variance from the additive genetic to the residual component (**files A2, A4**, electronic enhancements), which is not expected if *vgll3* comprises additive genetic effects on length.

Female condition heritability was, like that for female length and in contrast to that for maturation probability, similar between environments (h^2^CON_Warm_ - h^2^CON_Cold_ = −0.09, −0.30-0.08) and the between-environment genetic correlation of R_G_CON_Warm_._Cold_ = 0.83 (95% CI = 0.66-0.95) indicated very little genotype re-ranking between environments (**figure 4A-C**). Additive *vgll3* effects explained 15.0 and 6.0% of the condition heritability and 4.7 and 2.3% of the phenotypic variance in the warm and cold environments, respectively. And, *vgll3* effects explained some of the between-environment genetic correlation for female condition (**figure 4C; file A4**, electronic enhancement), although less than what *vgll3* explained for maturation of the same parameter. Thus, unlike for maturation, for which we detected stable *vgll3* effects that explained a large share of the genetic between-environment correlation in the presence of a significant heritability difference between environments, the environmentally stable *vgll3* condition effects were detected in the presence of a relatively more stable genetic background for condition.

We collected additional evidence for determining the phenotypic path how *vgll3* affects maturation timing by estimating between-trait genetic correlation between sexes. By quantifying the proportion of the additive *vgll3* effect contribution to trait-specific heritability, we were unable to support a hypothesized *vgll3* association for maturation via length but support an association for maturation via condition. Quantitative contribution estimates of the genetic marker to the heritability were sufficiently large to consider *vgll3* a large-effect locus for both maturation and condition. Nonetheless, *vgll3* contributions to the phenotypic maturation-timing variance were lower than previous estimates at a later life stage (Ayllon et al. 2015; Barson et al. 2015; Ayllon et al. 2019), agreeing with the assumption that initially detected genetic-marker effects tend to be overestimated (Hill 2010), and which could have several biological, methodological, or statistical reasons (discussion of which goes beyond the scope of this manuscript). Furthermore, by having observed the same direction for *vgll3* effects between male maturation and female condition, we had obtained the first indication that *vgll3*-induced variation for maturation and condition may underlie a common *vgll3*-related molecular process. We wanted to further foster this idea by quantifying the *vgll3* contribution to the genetic correlation between male maturation and female length (MAT.LEN) or female condition (MAT.CON), and thus estimated the between-trait genetic correlation between sexes.

We did not detect genetic correlations between male maturation and female (or even male) length, although estimates in both environments were positive and the relationship-controlled phenotypic correlation estimate (R_P_) (Searle 1961) between male maturation and male length was significant in both environments (**figure 4D, E**). Furthermore, all correlation estimates (genetic and phenotypic) between male maturation and female or male length increased after accounting for *vgll3* effects (**figure 4D, E**). Thus, we were able to further exclude length as a mediator of *vgll3* effects on maturation, which requires inferring a positive genetic covariance among *vgll3* effects between traits, but instead found support for a negative genetic covariance among *vgll3* effects between maturation and length.

Between male maturation and female (and male) condition, we, however, detected a positive genetic correlation, although only in the warm environment (we discuss this environmental difference below) (**figure 4D, E**). After accounting for *vgll3* effects, all genetic correlation estimates between male maturation and female (or male) condition decreased, as is expected under scenarios of either the presence of pleiotropic *vgll3* effects on condition and maturation, or when one trait mediates the genetic-marker associated variation of the other. The decreasing effect was strongest for the genetic correlation between male maturation and female condition, which even rendered non-significant when accounting for *vgll3* effects (**figure 4D**). Ignoring a *vgll3* pleiotropy scenario, this latter result suggests a positive genetic covariance among the significant *vgll3* effects between male maturation and female condition. This covariance may even be dominated by *vgll3* effects and, thereby, suggests a *vgll3* effect mediation on maturation via condition and not *vice versa* (because maturation variation cannot account for condition variation in immature females).

### *Vgll3* as a candidate gene for a resource allocation trade-off?

Although many effects for length were statistically non-significant, we cannot entirely rule out small *vgll3* effects on length. Generally, growth or length and maturation initiation are assumed to associate positively (Taranger et al. 2010; Andersson et al. 2018). By inferring a negative covariance among *vgll3* effects between maturation and length we were able to rule out length mediating *vgll3* effects on maturation with a high certainty. However, the inferred negative genetic covariance among *vgll3* effects between maturation and length support a scenario for a *vgll3*-governed resource allocation trade-off. Such a trade-off was also, albeit weakly, indicated by the non-significant 1% length decrease per *vgll3*\*E allele that was accompanied by a significant 1% condition increase. This result is typical in life-history research for traits that represent within-individual resource allocation trade-offs in the presence of relatively much larger among-individual variation for resource acquisition (de Jong and van Noordwijk 1992; Roff 2002). The idea of a *vgll3*-mediated resource allocation trade-off also makes biological sense. This is not only because resource allocation theory predicts that, regardless of underlying genetics, energy allocated to condition cannot concurrently be allocated to growth, but also due to the suggested role of Vgll3 in controlling mesenchymal cell fate into either adipocytes (thus increasing body condition) or bone and cartilage lineages (thus increasing somatic growth) (Halperin et al. 2013). However, a final conclusion for our tentative results requires additional research, also because there are other plausible candidate genes in the same genome region as *vgll3*, e.g., *akap11* (Ayllon et al. 2015; Barson et al. 2015).

### The presence of strong temperature effects on maturation timing and body condition

A remaining question is why we detected a significant genetic correlation between male maturation and female condition in the warm, but not the cold, environment. We suspect that one reasons may be that the higher maturation rate and maturation heritability in the warm provided sufficient information to detect the genetic correlation, because higher heritabilities decrease the error of the genetic correlation estimate (Robertson 1959). Furthermore, many males in the cold environment with “high” breeding values for condition (adjusted for temperature environment) did not mature, possibly because of the much lower average condition in the cold, or due to other lower temperature-related causes, such as not exceeding growth or size thresholds needed for maturation (Taranger et al. 2010; Andersson et al. 2018), masking an otherwise positive genetic correlation.

Extending these thoughts, we detected environmental effects on condition that were consistent with the notion that both condition and maturation covary positively with temperature. Specifically, when translated to the proportional scale, we detected a 4.6% (95% CI = 2.9-6.4%) higher female condition in the warm relative to the cold environment, where we had also detected a much higher maturation probability (0.41, 0.27-0.58 vs. 0.05, 0.02-0.08; **figure 3A, B, E, F**). Thus, the environmental effect of a global-warming-relevant 2 °C temperature difference (IPCC 2014) on condition (4.6%) was more than twice the maximum genetic *vgll3* effect (EE vs. LL: 2%) and may have contributed considerably to the large environmental temperature effect on maturation probability. Similar positive relationships between body condition and water temperature have previously been observed (Dwyer and Piper 1987; Jonsson et al. 2013; Tromp et al. 2018). We suspect that temperature effects on maturation via condition explain, at least partly, unexpectedly high maturation rates in heated salmon rearing facilities (Good and Davidson 2016) and why effects on size and maturation timing vary between growth acceleration through increase in feed vs. temperature (Berrigan and Charnov 1994; Shapiro Goldberg et al. 2019). Important for consistency of natural and human selection success across temperatures (Falconer 1952), the large average temperature difference for condition observed here was evident in the presence of both high genetic correlations (> 0.8 for both sexes) and similar female condition heritabilities between temperatures (**figure 4A-C**). Furthermore, *vgll3* effect estimates on both female condition and on male maturation did not differ between temperature environments, and *vgll3* effects contributed to the between-environment genetic correlations for both traits. Thus, the environmental stability of *vgll3* condition effects contributes to the general environmental stability of additive genetic condition effects.

These new results suggest - together with the previous results on maturation (Ayllon et al. 2015; Barson et al. 2015; Ayllon et al. 2019) - the presence of environmentally stable *vgll3* effects on both condition and maturation. A large share of *vgll3* effects for the positive genetic correlation between maturation and condition predicts rapid evolutionary co-responses to selection for either trait, but also predicts that their genetic correlation and importance is sensitive to *vgll3* allele frequencies. The sex-specific results, together with previous biological knowledge on their causal association (Taranger et al. 2010; Andersson et al. 2018), indicate that large *vgll3* effects on maturation are likely mediated via large condition effects and suggest *vgll3* as a candidate locus for controlling the resource allocation trade-off between energy reserves and somatic growth.

## Supporting information

electronic enhancements

## Literature Cited

Anderson, E. C. 2010. Computational algorithms and user-friendly software for parentage-based tagging of Pacific salmonids. Final report submitted to the Pacific Salmon Commission’s Chinook Technical Committee (US Section). Pages 46.

Andersson, E., G. L. Taranger, A. Wargelius, and R. W. Schulz. 2018. Puberty in Fish, Pages 426–429 in M. K. Skinner, ed. Encyclopedia of Reproduction. Oxford, Academic Press.

Aykanat, T., M. Lindqvist, V. L. Pritchard, and C. R. Primmer. 2016. From population genomics to conservation and management: a workflow for targeted analysis of markers identified using genome-wide approaches in Atlantic salmon *Salmo salar*. Journal of Fish Biology 89:2658–2679.

Ayllon, F., E. Kjaerner-Semb, T. Furmanek, V. Wennevik, M. F. Solberg, G. Dahle, G. L. Taranger et al. 2015. The *vgll3* locus controls age at maturity in wild and domesticated Atlantic salmon (*Salmo salar* L.) males. PLoS Genetics 11:e1005628.

Ayllon, F., M. F. Solberg, K. A. Glover, F. Mohammadi, E. Kjaerner-Semb, P. G. Fjelldal, E. Andersson et al. 2019. The influence of *vgll3* genotypes on sea age at maturity is altered in farmed mowi strain Atlantic salmon. BMC Genetics 20:44.

Barson, N. J., T. Aykanat, K. Hindar, M. Baranski, G. H. Bolstad, P. Fiske, C. Jacq et al. 2015. Sex-dependent dominance at a single locus maintains variation in age at maturity in salmon. Nature 528:405–408.

Bell, J. A., D. Carslake, K. H. Wade, R. C. Richmond, R. J. Langdon, E. E. Vincent, M. V. Holmes et al. 2018. Influence of puberty timing on adiposity and cardiometabolic traits: a Mendelian randomisation study. PLoS Medicine 15:e1002641.

Berrigan, D., and E. L. Charnov. 1994. Reaction norms for age and size at maturity in response to temperature: a puzzle for life historians. Oikos 70:474.

Butler, D. G., B. R. Cullis, A. R. Gilmour, and B. J. Gogel. 2009, Mixed models for S language environments ASReml-R reference manual. Brisbane, Australia, Queensland Department of Primary Industries and Fisheries, NSW Department of Primary Industries.

Cousminer, D. L., D. J. Berry, N. J. Timpson, W. Ang, E. Thiering, E. M. Byrne, H. R. Taal et al. 2013. Genome-wide association and longitudinal analyses reveal genetic loci linking pubertal height growth, pubertal timing and childhood adiposity. Human Molecular Genetics 22:2735–2747.

Czorlich, Y., T. Aykanat, J. Erkinaro, P. Orell, and C. R. Primmer. 2018. Rapid sex-specific evolution of age at maturity is shaped by genetic architecture in Atlantic salmon. Nature Ecology & Evolution 2:1800–1807.

Danielsen, E. T., M. E. Moeller, and K. F. Rewitz. 2013. Nutrient signaling and developmental timing of maturation. Current Topics in Developmental Biology 105:37–67.

de Jong, G., and A. J. van Noordwijk. 1992. Acquisition and allocation of resources - genetic (co)variances, selection, and life histories. American Naturalist 139:749–770.

de Villemereuil, P., O. Gimenez, B. Doligez, and R. Freckleton. 2013. Comparing parent-offspring regression with frequentist and Bayesian animal models to estimate heritability in wild populations: a simulation study for Gaussian and binary traits. Methods in Ecology and Evolution 4:260–275.

de Villemereuil, P., H. Schielzeth, S. Nakagawa, and M. Morrissey. 2016. General methods for evolutionary quantitative genetic inference from generalized mixed models. Genetics 204:1281–1294.

Dunlop, E. S., M. Heino, and U. Dieckmann. 2009. Eco-genetic modeling of contemporary life-history evolution. Ecological Applications 19:1815–1834.

Dupont, J., M. Reverchon, M. J. Bertoldo, and P. Froment. 2014. Nutritional signals and reproduction. Molecular and Cellular Endocrinology 382:527–537.

Dwyer, W. P., and R. G. Piper. 1987. Atlantic salmon growth efficiency as affected by temperature. The Progressive Fish-Culturist 49:57–59.

Enberg, K., C. Jørgensen, E. S. Dunlop, Ø. Varpe, D. S. Boukal, L. Baulier, S. Eliassen et al. 2012. Fishing-induced evolution of growth: concepts, mechanisms and the empirical evidence. Marine Ecology 33:1–25.

Falconer, D. S. 1952. The problem of environment and selection. American Naturalist 86:293–298.

Fleming, I. A. 1996. Reproductive strategies of Atlantic salmon: Ecology and evolution. Reviews in Fish Biology and Fisheries 6:379–416.

Gjedrem, T., and M. Baranski. 2005, Selective Breeding in Aquaculture: An Introduction: Reviews: Methods and Technologies in Fish Biology and Fisheries. London, Springer.

Good, C., and J. Davidson. 2016. A review of factors influencing maturation of Atlantic salmon, *Salmo salar*, with focus on water recirculation aquaculture system environments. Journal of the World Aquaculture Society 47:605–632.

Hadfield, J. D. 2010. MCMC methods for multi-response generalized linear mixed models: the MCMCglmm R package. Journal of Statistical Software 33:1–22.

Hadfield, J. D., and R. B. O’Hara. 2015. Increasing the efficiency of MCMC for hierarchical phylogenetic models of categorical traits using reduced mixed models. Methods in Ecology and Evolution 6:706–714.

Halperin, D. S., C. Pan, A. J. Lusis, and P. Tontonoz. 2013. Vestigial-like 3 is an inhibitor of adipocyte differentiation. Journal of Lipid Research 54:473–481.

Henderson, C. R. 1973. Sire evaluation and genetic trends. Journal of Animal Science 1973:10–41.

Hill, W. G. 2010. Understanding and using quantitative genetic variation. Philosophical Transactions of the Royal Society B: Biological Sciences 365:73–85.

IPCC. 2014. Future Climate Changes, Risk and Impacts, Pages 56–74 in Core Writing Team, R. K. Pachauri, and L. A. Meyer, eds. Climate Change 2014: Synthesis Report. Contribution of Working Groups I, II and III to the Fifth Assessment Report of the Intergovernmental Panel on Climate Change. Geneva, Switzerland, IPCC.

Jones, O. R., and J. Wang. 2010. COLONY: a program for parentage and sibship inference from multilocus genotype data. Molecular Ecology Resources 10:551–555.

Jonsson, B., N. Jonsson, and A. G. Finstad. 2013. Effects of temperature and food quality on age and size at maturity in ectotherms: an experimental test with Atlantic salmon. Journal of Animal Ecology 82:201–210.

Kardos, M., and G. Luikart. 2019. The genetic architecture of fitness drives population viability during rapid environmental change. BioRxiv. doi: 10.1101/660803v2.

Kause, A., O. Ritola, T. Paananen, E. Mäntysaari, and U. Eskelinen. 2003. Selection against early maturity in large rainbow trout *Oncorhynchus mykiss*: the quantitative genetics of sexual dimorphism and genotype-by-environment interactions. Aquaculture 228:53–68.

Kenward, M. G., and J. H. Roger. 1997. Small sample inference for fixed effects from restricted maximum likelihood. Biometrics 53:983–997.

Kuparinen, A., and J. A. Hutchings. 2017. Genetic architecture of age at maturity can generate divergent and disruptive harvest-induced evolution. Philosophical Transactions of the Royal Society of London. Series B: Biological Sciences 372.

Lande, R. 1982. A quantitative genetic theory of life-history evolution. Ecology 63:607–615.

Law, R. 2007. Fisheries-induced evolution: present status and future directions. Marine Ecology Progress Series 335:271–277.

Lynch, M., and B. Walsh. 1998, Genetics and Analysis of Quantitative Traits. Sunderland, Massachusetts, Sinauer.

Marschall, E. A., T. P. Quinn, D. A. Roff, J. A. Hutchings, N. B. Metcalfe, T. A. Bakke, R. L. Saunders et al. 1998. A framework for understanding Atlantic salmon (*Salmo salar*) life history. Canadian Journal of Fisheries and Aquatic Sciences 55:48–58.

Meerburg, D. J. 1986, Salmonid age at maturity: Canadian Special Publication of Fisheries and Aquatic Sciences, v. 89. Ottawa, Department of Fisheries and Oceans.

Nakayama, K., J. Ohashi, K. Watanabe, L. Munkhtulga, and S. Iwamoto. 2017. Evidence for very recent positive selection in Mongolians. Molecular Biology and Evolution 34:1936–1946.

Robertson, A. 1959. The sampling variance of the genetic correlation coefficient. Biometrics 15:469–485.

Roff, D. A. 2002, Life History Evolution. Sunderland, Massachusetts, Sinauer Associates, Inc.

Rowe, D. K., J. E. Thorpe, and A. M. Shanks. 1991. Role of fat stores in the maturation of male Atlantic salmon (*Salmo salar*) parr. Canadian Journal of Fisheries and Aquatic Sciences 48:405–413.

Sae-Lim, P., B. Gjerde, H. M. Nielsen, H. Mulder, and A. Kause. 2016. A review of genotype-by-environment interaction and micro-environmental sensitivity in aquaculture species. Reviews in Aquaculture 8:369–393.

Searle, S. R. 1961. Phenotypic, genetic and environmental correlations. Biometrics 17:474–480.

Shapiro Goldberg, D., I. van Rijn, M. Kiflawi, J. Belmaker, and M. Heino. 2019. Decreases in length at maturation of Mediterranean fishes associated with higher sea temperatures. ICES Journal of Marine Science 76:946–959.

Stearns, S. C. 1992, The Evolution of Life Histories. New York, Oxford University Press.

Stearns, S. C., and J. C. Koella. 1986. The evolution of phenotypic plasticity in life-history traits: predictions of reaction norms for age and size at maturity. Evolution 40:893–913.

Stram, D. O., and J. W. Lee. 1994. Variance components testing in the longitudinal mixed effects model. Biometrics 50:1171–1177.

Sutton, S. G., T. P. Bult, and R. L. Haedrich. 2000. Relationships among fat weight, body weight, water weight, and condition factors in wild Atlantic salmon parr. Transactions of the American Fisheries Society 129:527–538.

Taranger, G. L., M. Carrillo, R. W. Schulz, P. Fontaine, S. Zanuy, A. Felip, F. A. Weltzien et al. 2010. Control of puberty in farmed fish. General and Comparative Endocrinology 165:483–515.

Trombley, S., A. Mustafa, and M. Schmitz. 2014. Regulation of the seasonal leptin and leptin receptor expression profile during early sexual maturation and feed restriction in male Atlantic salmon, *Salmo salar* L., parr. General and Comparative Endocrinology 204:60–70.

Tromp, J. J., P. L. Jones, M. S. Brown, J. A. Donald, P. A. Biro, and L. O. B. Afonso. 2018. Chronic exposure to increased water temperature reveals few impacts on stress physiology and growth responses in juvenile Atlantic salmon. Aquaculture 495:196–204.

Tu, W., E. K. Wagner, G. J. Eckert, Z. Yu, T. Hannon, J. H. Pratt, and C. He. 2015. Associations between menarche-related genetic variants and pubertal growth in male and female adolescents. Journal of Adolescent Health 56:66–72.

Wells, J. C. K., R. M. Nesse, R. Sear, R. A. Johnstone, and S. C. Stearns. 2017. Evolutionary public health: introducing the concept. The Lancet 390:500–509.

