## appendix for "Large single-locus effects for maturation timing are mediated via body condition in Atlantic salmon"

- Settings for the parental reconstruction
- Figure A1: Non-detected effects for maturation probability.
- Figure A2: Comparison between univariate or bivariate with multivariate model estimates.
- Figure A3: Temperature-by-additive *vgll3* effects were not detected.
- Table A1: Maternal effects were not detected for body length or body condition.

### Settings for the parental reconstruction

We reconstructed the grandparents of the experimental cohort by using 131 SNPs genotyped for 984 potential parents of the experimental cohort in COLONY 2.0.6.5 (Jones and Wang, 2010). As settings other than the default, we used the outbreeding model, under a polygamy mating mode, updated allele frequencies by accounting for the inferred relationship, invoked a complexity prior, chose one long runs under the Full-Likelihood method with high precision. Genotyping error rates remained unknown and we therefore defined them as equal among loci with a rate of 0.001.

Jones, O. R., & Wang, J. (2010). COLONY: a program for parentage and sibship inference from multilocus genotype data. Mol. Ecol. Resour. 10: 551-555.

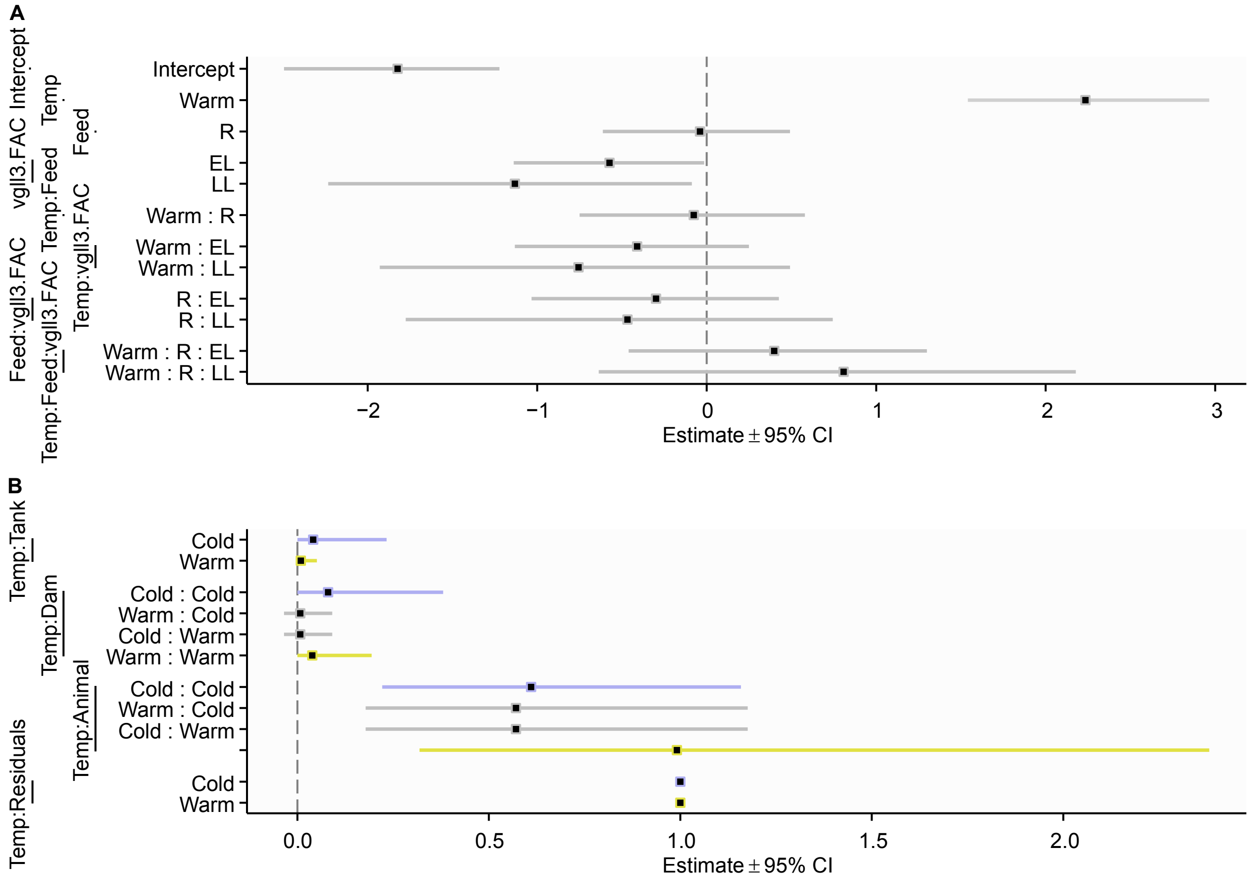


**Figure A1:** Non-detected effects for maturation probability.

(**A**) Mean and contrast effect estimates (grouped by term, with term level indicated) with 95% credible intervals for the univariate threshold model on male maturation binaries with initial mean effect structure. Maturation binaries were modelled as a linear function of *vgll3* genotype (vgll3.FAC; levels: EE, EL, LL), temperature treatment (Temp; levels: Cold, Warm), feed treatment (Feed, levels: R = temporally restricted, F = full feeding), and all interactions. All feed treatment effects were removed for a reduced model because their 95% credible intervals included zero. (**B**) Variance and covariance term estimates (grouped by term, with term level indicated) for the univariate threshold model on male maturation binaries with the reduced final mean effect structure. Maturation binaries were modelled as a linear function of *vgll3* genotype (vgll3.FAC; levels: EE, EL, LL) and temperature (Temp; levels: Cold, Warm). This model included (co)variances for dam effects (Dam) that were removed for the final model because their 95% credible intervals included zero. Variance terms for tank effects (Tank) were retained, although their 95% credible intervals included zero, because tank effects reflect the experimental design. Residual variance in each environment was fixed at one to make additive genetic (animal) effect variance estimable. As a result, all (co)variance estimates are scaled relative to a residual variance of one.

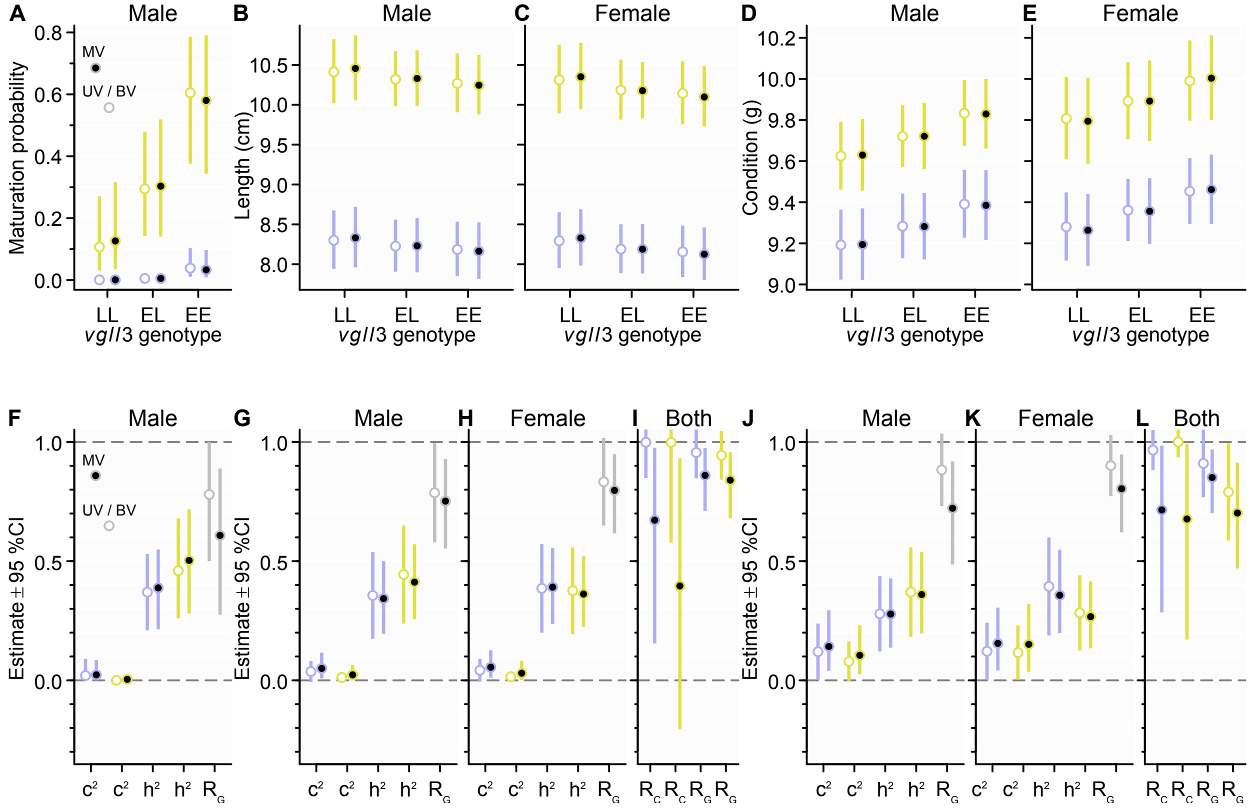


**Figure A2.** Comparison between univariate or bivariate with multivariate model estimates.

(**A, F**) Bayesian MCMC univariate (UV) and multivariate (MV) model estimates for maturation probability with posterior-based 95% credible intervals. (**B, C, G, H, I**) REML bivariate (BV) model estimates with Taylor-series-derived approximate 95% confidence intervals (partly off the plot area for variance proportions or correlations > 1) and Bayesian MCMC multivariate (MV) model estimates with posterior-based 95% credible intervals for sex-specific body length. (**D, E, J, K, L**) REML bivariate (BV) model estimates with Taylor-series-derived approximate 95% confidence intervals (partly off the plot area for variance proportions or correlations > 1) and Bayesian MCMC multivariate (MV) model estimates with posterior-based 95% credible intervals for sex-specific body condition. Variance component proportions (**F, G, H, J, K**) are for common environmental variance ($c^2$; based on tank replicates) or additive genetic variance ($h^2$; based on expected relatedness among individuals), whereas correlations are for additive genetic correlation ($R_G$) between temperature environments (**G, H, J, K**) or for common environmental correlation ($R_C$) and additive genetic correlation ($R_G$) between sexes within environments (**I, L**). Estimates in all panels for the warm environment are in yellow, whereas estimates for the cold environment are in blue. Gray colour indicates between-temperature environment correlations.

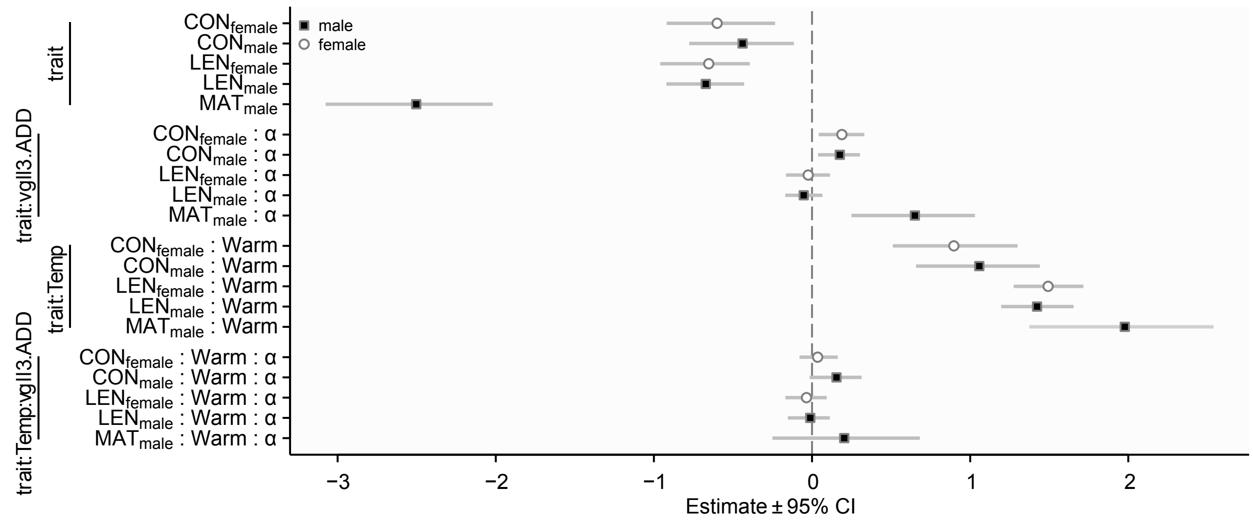


**Figure A3:** Temperature-by-additive *vgll3* effects were not detected.

Mean and contrast effect estimates (grouped by term, with term level indicated) with 95% credible intervals for the multivariate animal model on traits of male maturation ($MAT_{male}$), male length ($LEN_{male}$), female length ($LEN_{female}$), male condition ($CON_{male}$), and female condition ($CON_{female}$). Traits were modelled as a linear function of temperature effects (Temp; levels: Cold, Warm), additive *vgll3* effects (vgll3.ADD; for EE, EL, LL; α = 1, 0, -1), and temperature-by-additive *vgll3* effects (Temp:vgll3.ADD). Temperature-by-additive *vgll3* effects for all traits were removed for the final model because their 95% credible intervals included zero.

**Table A1:** Maternal effects were not detected for body length or body condition. Results by likelihood ratio tests (LRT) on the covariance structure in bivariate models for the traits of either male and female body length (LEN) or male and female body condition (CON). K indicates the total number of covariance parameters in the model, Loglik is the log of the model likelihood, and AIC is the Akaike information criterion with a "smaller is better" definition. The model # in bold is the chosen model.

| Model # | Dam covariance matrix | Description | K | Loglik | AIC | vs. # | ΔAIC | LRT | *P* |
|---|---|---|---|---|---|---|---|---|---|
| **LEN** | | | | | | | | | |
| 0 | Trait:Temp:Dam | unstructured 4x4 | 32 | -335.2 | 734.4 | - | - | - | - |
| 1 | Trait:Dam | unstructured 2x2 | 25 | -335.0 | 719.9 | 0 | 14.4 | -0.4 | 1.00 |
| 2 | Temp:Dam | unstructured 2x2 | 25 | -335.1 | 720.1 | 0 | 14.2 | -0.2 | 1.00 |
| 3 | Dam | common variance | 23 | -335.1 | 718.0 | 1 | 3.6 | 0.4 | 0.833 |
| 3 | Dam | common variance | 23 | -335.1 | 718.0 | 2 | 3.8 | 0.2 | 0.914 |
| **4** | NA | no dam effects | 22 | -335.5 | 2523.7 | 3 | 1.3 | 0.7 | 0.409 |
| **CON** | | | | | | | | | |
| 0 | Trait:Temp:Dam | unstructured 4x4 | 32 | -1237.1 | 2538.3 | - | - | - | - |
| 1 | Trait:Dam | unstructured 2x2 | 25 | -1238.8 | 2527.5 | 0 | 10.8 | 3.3 | 0.861 |
| 2 | Temp:Dam | unstructured 2x2 | 25 | -1238.0 | 2526.0 | 0 | 12.3 | 1.7 | 0.973 |
| 3 | Dam | common variance | 23 | -1238.8 | 2523.7 | 1 | 3.9 | 0.1 | 0.936 |
| 3 | Dam | common variance | 23 | -1238.8 | 2523.7 | 2 | 3.9 | 1.7 | 0.437 |
| **4** | NA | no dam effects | 22 | -1239.7 | 2523.3 | 3 | 0.3 | 1.7 | 0.196 |
